# Single-cell transcriptomics reveal individual and synergistic effects of Trisomy 21 and GATA1s on hematopoiesis

**DOI:** 10.1101/2024.05.24.595827

**Authors:** Kaoru Takasaki, Eric K. Wafula, Sara S. Kumar, David Smith, Ying Ting Sit, Alyssa L. Gagne, Deborah L. French, Christopher S. Thom, Stella T. Chou

**Author notes:** **Corresponding Author**: Stella T. Chou, 3615 Civic Center Boulevard, Abramson Research Center Room 316D, Philadelphia, PA 19104.

## Abstract

Trisomy 21 (T21), or Down syndrome (DS), is associated with baseline macrocytic erythrocytosis, thrombocytopenia, and neutrophilia, as well as transient abnormal myelopoiesis (TAM) and myeloid leukemia of DS (ML-DS). TAM and ML-DS blasts both arise from an aberrant megakaryocyte-erythroid progenitor and exclusively express GATA1s, the truncated isoform of *GATA1*, while germline *GATA1s* mutations in a non-T21 context lead to congenital cytopenia(s) without a leukemic predisposition. This suggests that T21 and GATA1s both perturb hematopoiesis in multipotent progenitors, but studying their individual effects is challenging due to limited access to relevant human progenitor populations. To dissect individual developmental impacts, we used single-cell RNA-sequencing to interrogate hematopoietic progenitor cells (HPCs) from isogenic human induced pluripotent stem cells differing only by chromosome 21 and/or *GATA1* status. The transcriptomes of these HPCs revealed significant heterogeneity and lineage skew dictated by T21 and/or GATA1s. T21 and GATA1s each disrupted temporal regulation of lineage-specific transcriptional programs and specifically perturbed cell cycle genes. Trajectory inference revealed that GATA1s nearly eliminated erythropoiesis, slowed MK maturation, and promoted myelopoiesis in the euploid context, while in T21 cells, GATA1s competed with the enhanced erythropoiesis and impaired megakaryopoiesis driven by T21 to promote production of immature erythrocytes, MKs, and myeloid cells. The use of isogenic cells revealed distinct transcriptional programs that can be attributed specifically to T21 and GATA1s, and how they independently and synergistically result in HPC proliferation at the expense of maturation, consistent with a pro-leukemic phenotype.

## Introduction

Trisomy 21 (T21), or Down syndrome (DS), is associated with macrocytic erythrocytosis, thrombocytopenia, and neutrophilia particularly in the newborn period,^1–4^ and transient abnormal myelopoiesis (TAM) and myeloid leukemia of DS (ML-DS).^5,6^ TAM and ML-DS blasts are characterized by somatic mutations in the essential hematopoietic transcription factor *GATA1* that result in exclusive expression of the amino (N) terminus-truncated isoform GATA1s^7–10^ with impaired transactivation activity.^7,11–14^ Both myeloproliferative disorders are attributed to an aberrant megakaryocyte-erythroid progenitor (MEP),^15–18^ and the blasts typically carry a megakaryocyte (MK)-like surface marker signature.

How T21 and GATA1s exert their hematopoietic phenotypes remains incompletely understood. Germline *GATA1s* mutations without T21 lead to congenital cytopenia(s) without leukemic predisposition,^19–21^ while leukemic clones with *GATA1s* mutations and acquired T21 have a ML-DS-like phenotype.^22–25^ Previous studies have shown that the extra copy of chromosome 21 (HSA21) perturbs gene expression and histone modifications throughout the genome, and that addition of a *GATA1s* mutation further perturbs acetylation specifically at genes involved in hematopoiesis and cell cycle regulation.^26–29^ These findings suggest that genome-wide gene dysregulation underlies the manifestations of DS and emphasize the unique interactions between T21 and *GATA1*.

Murine DS models only partially recapitulate human disease: HSA21 orthologs map to three murine chromosomes,^30^ DS mice tend to exhibit macrocytic anemia,^31,32^ and crosses between DS and *Gata1s* mice do not develop leukemia.^31,33,34^ Studies in human cells are limited by access to primary hematopoietic progenitor cells (HPCs) and individual variation in genetic backgrounds that overwhelm the transcriptional changes resulting from T21 and/or GATA1s,^35,36^ requiring isogenic or large-scale approaches.^27,28^ These challenges underscore the value of human cell model systems in which HSA21 or *GATA1* can be manipulated in isolation.

Human T21 induced pluripotent stem cell (iPSC) models accurately reflect DS hematopoietic phenotypes: T21 increases erythropoiesis, while GATA1s severely impairs erythropoiesis but enhances MK and myeloid proliferation in both euploid and T21 backgrounds.^37–42^ However, bulk assessments obscure the heterogeneity of HPCs,^43,44^ including those derived from iPSCs.^38,45,46^ Thus, to further dissect the developmental impacts of GATA1s on hematopoiesis in euploid and T21 cells, we performed single-cell RNA sequencing (scRNA-seq) of multipotent and lineage-committed HPCs derived from isogenic human iPSCs differing only by HSA21 copy number and/or GATA1 status. We examined transcriptome states at early and late HPC stages, which reveal distinct dynamics of lineage priming and fate decisions driven individually and synergistically by HSA21 and GATA1 status. In euploid cells, GATA1s led to a transcriptional state that nearly eliminated erythropoiesis, slowed MK maturation, and promoted myelopoiesis, while in T21 cells, GATA1s competed with the enhanced erythropoiesis and impaired megakaryopoiesis driven by T21 to give rise to immature erythrocytes, MKs, and myeloid cells, consistent with a myeloproliferative phenotype.

## Methods

### Generation of isogenic iPSCs

Isogenic T21/GATA1s,^47^ T21/wild-type GATA1 (wtGATA1),^47^ and euploid/GATA1s^48^ iPSCs were derived as previously described^49,50^ from an individual who presented with TAM and then in remission. The isogenic euploid/wtGATA1 clone was generated from the euploid/GATA1s clone with the CRISPR/Cas9 nuclease system and gRNAs (Integrated DNA Technologies) targeting the individual’s known *GATA1* mutation (**Figure 1A**). All iPSC lines were characterized by CNV (copy number variation) analysis, karyotype, *GATA1* Sanger sequencing (GENEWIZ, Inc.), and Western blot for GATA1 (**Supplemental Methods; Supplemental Figure 1**). Sequences of gRNAs, oligonucleotides for homologous recombination, and PCR and sequencing primers (Integrated DNA Technologies) are shown in **Supplemental Table 1**.

**Figure 1.**
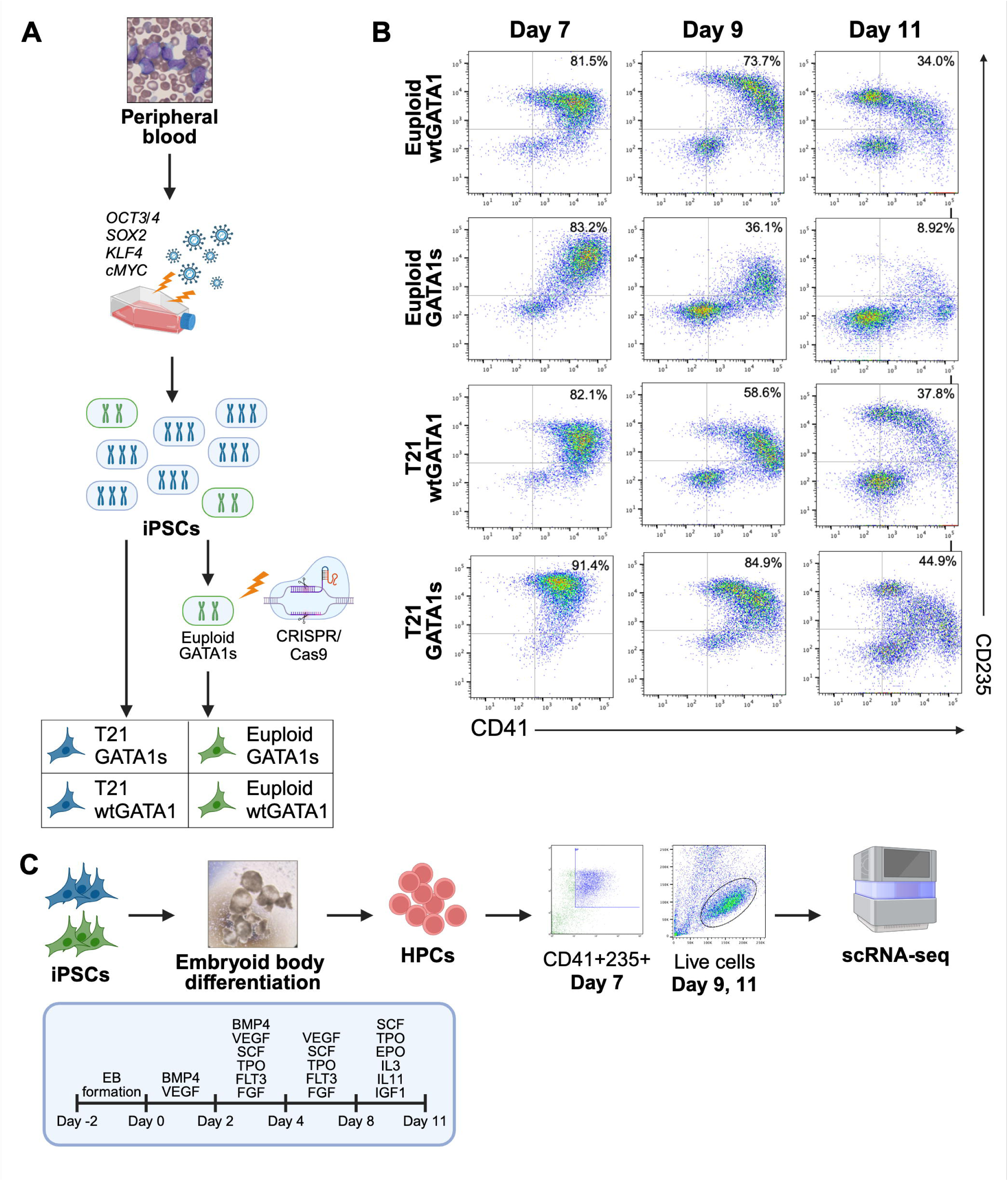
Characterization of HPCs differentiated from isogenic iPSCs. A) Generation of isogenic iPSCs differing by HSA21 and/or GATA1 status. B) Flow cytometry plots of Day 7, 9, and 11 HPCs differing only by HSA21 and/or GATA1 status. CD41^+^/CD235^+^ indicates multipotency, CD41^+^/CD235^-^ suggests megakaryocytic bias, CD41^-^/CD235^+^ suggests erythroid bias, and CD41^-^/CD235^-^ suggests myeloid bias. C) Differentiation, isolation, and characterization of Day 7, 9, and 11 HPCs.

### Hematopoietic differentiation of iPSCs

iPSCs were cultured and differentiated by embryoid body (EB) formation using modifications on a protocol previously described by our laboratory (**Figure 1C**).^37,38,47,48^ Details are described in **Supplemental Methods**.

### Isolation of HPCs

On Day 7 of differentiation, HPCs were collected from disaggregated EBs and the supernatant, and sorted for CD41^+^/CD235^+^ cells on a FACSAria Fusion (BD Biosciences) or MoFlo Astrios EQ (Beckman Coulter) (**Figure 1C**). On Days 9 and 11 of differentiation, late HPCs were collected from the supernatant only, with sorting for live cells (**Figure 1C**). Details are described in **Supplemental Methods**.

### Flow cytometry

Flow cytometry was performed on a FACSCanto II (BD Biosciences) as previously described,^38^ using conjugated antibodies for surface-marker staining (**Supplemental Table 3**). Data were analyzed using FlowJo Software (BD Biosciences).

### scRNA-seq

Sorted HPCs were resuspended in 0.04% BSA in PBS at 1000 cells/μL, and 2 x 10^4^ cells per sample were loaded onto gel beads-in-emulsion wells (10x Genomics). cDNA libraries were reverse-transcribed from extracted RNA using the Chromium Single Cell Platform (10x Genomics). cDNA quality and integrity were measured by TapeStation High Sensitivity DNA analysis (Agilent) before sequencing with a standard 3’ assay with the NovaSeq 6000 platform and S1 Reagent Kit v1.5 (Illumina).

### Single-cell profiling data analysis

Raw base calls (BCL) files were demultiplexed into FASTQ files using CellRanger (v7.1.0)^51^ *mkfastq*, then processed with *counts* to generate feature-barcode counts matrices aligned to the 10x Genomics Human reference GRCh38 (Ensembl98). Contaminating ambient RNA expression was corrected using SoupX (v1.6.2)^64^ and Doublets (DoubletFinder v2.0.3),^53^ and cells with high mitochondrial RNA (**Supplemental Table 4**) were removed. High-quality data representing all differentiation time points were merged and individual sample objects normalized (*sctransform*) with Seurat (v4.4.0).^46^ Samples were integrated using the top 6000 highly variable genes and first 20 principal components. Cells were clustered using the Louvain algorithm (resolution 0.4).

Clusters were manually annotated based on calculated module scores for lineage-specific expression of selected hematopoietic marker genes (**Supplemental Table 5**) and previously published gene sets^34,47,48^ (Seurat *AddModuleScore*). Differential gene expression was assessed (Seurat *FindMarkers*) for cells with the same identity within each time point, comparing integrated GATA1s and wtGATA1 samples. Genes detected in ≥25% of cells in the two samples, with log-fold change (log_2_FC) >0.25 and adjusted *p*-value <0.05, were considered significantly differentially expressed. Gene set enrichment analysis (GSEA) implemented through fgsea v1.30.0^54,55^ was tested against the MsigDB Hallmark collection (v7.5.1).^54,56^ To simplify interpretation and assess the effects of GATA1s in euploid vs. T21 contexts, comparisons were restricted to within HSA21 status.

For RNA velocity analysis, CellRanger bam files were used to count spliced and unspliced transcripts (velocyto v0.17.17).^57^ RNA velocities and cell latent times were estimated with scVelo (v0.3.1).^58^ Pre-processed data from Seurat and velocyto-derived expression matrices were merged, then filtered and normalized using the top 6000 highly variable genes with at least 20 counts per gene in both spliced and unspliced matrices. Each cell’s first– and second-order moments were computed with the default parameters and velocities were estimated using the dynamical mode.

Cell differentiation trajectories were inferred using the CytoTRACE^59^ kernel of CellRank (v2.0.2).^60^ CytoTRACE scores and pseudotimes were computed on the previously-generated scVelo count matrices. Transition probabilities were calculated, and the first four predicted macrostates, which neatly matched annotated cell lineages, were set as the target terminal states (**Figure 4C; Supplemental Figure 7B**). Fate probabilities for cells to transition to the terminal states were estimated and driver genes were determined.

Genes that promote stalled lineage maturation in T21/GATA1s cells were identified by fitting a regression model testing the effect of GATA1s on the ploidy conditions while treating the estimated latent times as a continuous covariate (i.e. GATA1*T21 + latent time) (MAST v1.30.0).^61^

### Statistical analysis

Continuous variables were assessed using the Wilcoxon rank-sum test and categorical data with the χ² test. A 2-tailed or adjusted *p*-value <0.05 was considered statistically significant.

### Study approval

All patients or legal guardians provided written informed consent to participate in this study. The CHOP Institutional Review Board approved the study protocol.

## Results

### HPCs demonstrate lineage bias in multi-cytokine media

Isogenic iPSCs differing only by HSA21 and/or GATA1 status were generated from an individual with DS (**Figure 1A**) and underwent hematopoietic differentiation. By flow cytometry, >80% of Day 7 HPCs from each genotype (euploid/wtGATA1, euploid/GATA1s, T21/wtGATA1, T21/GATA1s) were CD41^+^/CD235^+^, which are considered multipotent since they yield erythroid, myeloid, mixed, and MK colonies on methylcellulose assays.^37^ Yet each genotype had distinct profiles that suggested varying degrees of erythroid (CD235^+^) or MK (CD41^+^) lineage bias (**Figure 1B left**). On Days 9 and 11, with continued exposure to multi-cytokine media, HPCs demonstrated erythroid (CD41^-^/235^+^), MK (CD41^+^/CD235^-^), or myeloid (CD41^-^/CD235^-^) commitment (**Figure 1B middle and right; Supplemental Figure 2**). By Day 11, euploid/GATA1s HPCs completely lacked a CD41^-^/CD235^+^ erythroid population, consistent with the congenital anemia associated with exclusive expression of GATA1s.^19,21^ In contrast, a subpopulation of T21/GATA1s HPCs surprisingly remained CD235^+^ throughout.

To identify the transcriptional programs that govern lineage fate in the presence of T21 and/or exclusive expression of GATA1s, we performed scRNA-seq on flow-purified Day 7 CD41^+^/CD235^+^ and Day 9 and 11 live HPCs (**Figure 1C**). We obtained 148,750 HPCs with high-quality transcriptome profiles. Integration and cluster annotation of Day 7, 9, and 11 HPCs from all 4 isogenic lines identified 7 major cell clusters: unbiased HPCs, MK (2)-or erythroid-biased HPCs, MKs, erythroid cells, and myeloid cells (**Figure 2A-B; Supplemental Figures 3-6**). Expression of specific marker genes revealed clear transcriptional priming of erythroid, megakaryocytic, and myeloid lineages for each genotype, even in flow-purified CD41^+^/CD235^+^ HPCs (**Figure 2C-D**), underscoring the significant heterogeneity that was made evident in an isogenic background and only by single-cell approaches. The single-cell transcriptomes revealed the evolution of lineage-committed subpopulations over time (**Figure 2C-D**).

**Figure 2.**
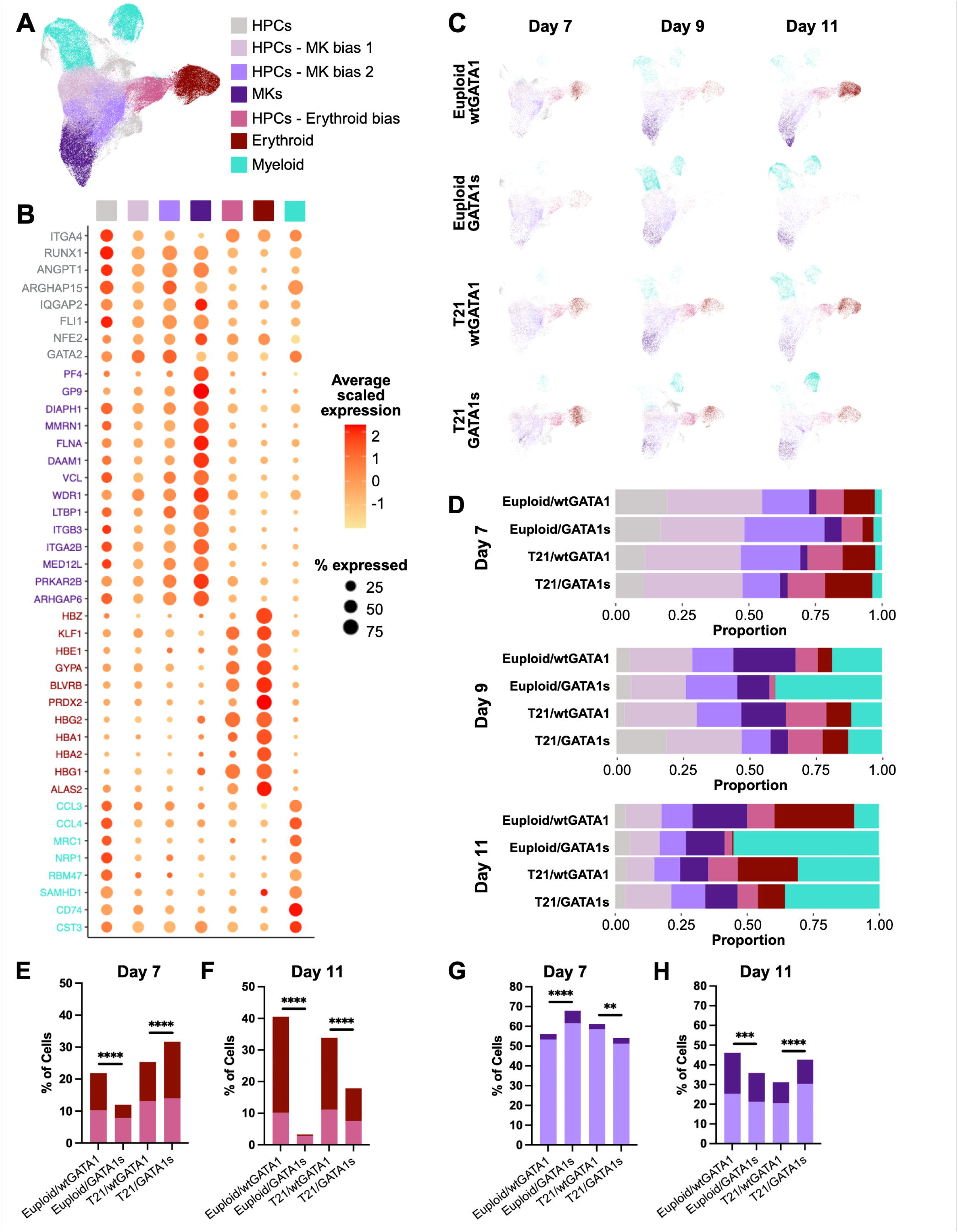
scRNA-seq characterization of HPCs generated from isogenic iPSCs differing by HSA21 and/or GATA1 status. A) Annotated integrated UMAP (Uniform Manifold Approximation and Projection) of euploid/wtGATA1, euploid/GATA1s, T21/wtGATA1, and T21/GATA1s HPCs collected on Days 7, 9, and 11. B) Relative expression of canonical hematopoietic genes, organized by lineage/cluster. C) Individual annotated UMAPs for each genotype and timepoint. * indicates distinct population of committed MKs in euploid/wtGATA1 cells. D) Relative proportions of cells in each lineage per genotype and timepoint. E) Percentage of erythroid-biased HPCs and committed erythroid cells within each genotype on Day 7. F) Percentage of erythroid-biased HPCs and committed erythroid cells within each genotype on Day 11. G) Percentage of MK-biased HPCs and committed MKs within each genotype on Day 7. H) Percentage of MK-biased HPCs and committed MKs within each genotype on Day 11. *p-*values reflect comparisons of relative proportions of biased HPCs vs. committed cells; \*\*\*\**p* <0.0001, \*\*\**p* = 0.0007, \*\**p* = 0.016.

### GATA1s impairs erythroid lineage commitment in euploid cells but results in an incomplete erythroid block in T21 cells

On Day 7, 21.9% of euploid/wtGATA1 cells were erythroid-biased HPCs or erythroid compared to 12.0% of euploid/GATA1s (*p* <0.0001) (**Figure 2D**; **Table 1**), consistent with the anemia observed with germline *GATA1s* mutations. T21 was associated with an erythroid drive, consistent with DS-associated polycythemia, but surprisingly T21/GATA1s cells exhibited the strongest erythroid skew at this stage (T21/wtGATA1: 25.4% vs. T21/GATA1s: 31.7%, *p* <0.0001) along with a greater proportion of committed, rather than erythroid-biased, cells (**Figure 2E**; **Table 1**).

**Table 1.** Frequencies of cells assigned to hematopoietic lineages in integrated analysis of Days 7-11 of differentiation, by genotype. Boxed sections highlight key findings.

|  |  | HPCs | HPCs –<br>Ery<br>bias | Ery | HPCs –<br>MK<br>bias 1 | HPCs –<br>MK<br>bias 2 | MK | Myeloid |
| --- | --- | --- | --- | --- | --- | --- | --- | --- |
| Day 7 | Euploid/wtGATA1 | 2517<br>(19.4%) | 1320<br>(10.2%) | 1515<br>(11.7%) | 2289<br>(17.7%) | 4615<br>(35.6%) | 362<br>(2.8%) | 336<br>(2.6%) |
|  | Euploid/GATA1s | 1544<br>(17.0%) | 709<br>(7.8%) | 381<br>(4.2%) | 2722<br>(30.0%) | 2843<br>(31.4%) | 582<br>(6.4%) | 278<br>(3.1%) |
|  | T21/wtGATA1 | 1209<br>(10.9%) | 1458<br>(13.1%) | 1367<br>(12.3%) | 2490<br>(22.4%) | 4014<br>(36.1%) | 306<br>(2.7%) | 284<br>(2.6%) |
|  | T21/GATA1s | 1322<br>(10.4%) | 1774<br>(14.0%) | 2243<br>(17.7%) | 1776<br>(14.0%) | 4708<br>(37.2%) | 369<br>(2.9%) | 463<br>(3.7%) |
| Day 9 | Euploid/wtGATA1 | 591<br>(5.1%) | 970<br>(8.3%) | 635<br>(5.4%) | 1801<br>(15.4%) | 2747<br>(23.5%) | 2747<br>(23.5%) | 2193<br>(18.8%) |
|  | Euploid/GATA1s | 789<br>(5.6%) | 285<br>(2.0%) | 21<br>(0.1%) | 2742<br>(19.3%) | 2915<br>(20.6%) | 1736<br>(12.2%) | 5691<br>(40.1%) |
|  | T21/wtGATA1 | 389<br>(3.0%) | 1930<br>(15.1%) | 1206<br>(9.4%) | 2147<br>(16.8%) | 3474<br>(27.2%) | 2159<br>(16.9%) | 1475<br>(11.5%) |
|  | T21/GATA1s | 2854<br>(18.9%) | 1963<br>(13.0%) | 1467<br>(9.7%) | 1636<br>(10.8%) | 4291<br>(28.4%) | 1001<br>(6.6%) | 1921<br>(12.7%) |
| Day 11 | Euploid/wtGATA1 | 625<br>(4.1%) | 1562<br>(10.2%) | 4625<br>(30.3%) | 1784<br>(11.7%) | 2081<br>(13.6%) | 3176<br>(20.8%) | 1422<br>(9.3%) |
|  | Euploid/GATA1s | 554<br>(5.7%) | 270<br>(2.8%) | 48<br>(0.5%) | 963<br>(9.9%) | 1093<br>(11.3%) | 1430<br>(14.7%) | 5340<br>(55.1%) |
|  | T21/wtGATA1 | 500<br>(4.2%) | 1336<br>(11.1%) | 2736<br>(22.8%) | 1179<br>(9.8%) | 1268<br>(10.6%) | 1289<br>(10.7%) | 3704<br>(30.8%) |
|  | T21/GATA1s | 457<br>(3.7%) | 928<br>(7.6%) | 1255<br>(10.3%) | 1558<br>(12.8%) | 2131<br>(17.5%) | 1500<br>(12.3%) | 4364<br>(35.8%) |

Ery = erythroid

The suppressive effect of GATA1s on erythropoiesis became more apparent by Day 11. Committed erythroid cells were the most abundant in the euploid/wtGATA1 sample (30.3%) but negligible in the euploid/GATA1s (0.5%) (*p* <0.0001) (**Figure 2F**; **Table 1**). The early erythroid drive in T21/GATA1s cells was no longer as strong: compared to T21/wtGATA1, there were fewer committed erythroid cells and proportionally more erythroid-biased HPCs, implying a block in lineage commitment that manifests at later points of hematopoietic development (*p* <0.0001 for both comparisons) (**Figure 2D,F**; **Table 1**). These findings suggest impaired, but not negligible, erythropoiesis resulting from the competing effects of T21 and GATA1s.

### GATA1s enhances megakaryopoiesis in euploid and T21 cells at distinct developmental timepoints

On Day 7, MK-biased HPCs and MKs together represented the largest subgroup of each genotype (**Figure 2D**; **Table 1**). GATA1s enhanced megakaryocytic lineage skew in euploid cells (euploid/wtGATA1: 56.1% vs. euploid/GATA1s: 67.9%, *p* <0.0001) and furthermore appeared to promote early commitment, with more of these cells representing MKs rather than MK-biased HPCs (euploid/wtGATA1 5.0% vs. euploid/GATA1s 9.5%, *p* <0.0001) (**Figure 2G**). However, these trends were reversed by Day 11. Of the 4 genotypes, only euploid/wtGATA1 developed a distinct population of committed MKs (**Figure 2C asterisk**). The euploid/GATA1s sample contained fewer megakaryocytic cells overall (euploid/wtGATA1: 46.1% vs. euploid/GATA1s: 35.9%, *p* <0.0001) and proportionately fewer committed MKs compared to euploid/wtGATA1 (**Figure 2H**). By Day 11, megakaryocytic cells represented the largest subpopulation of T21/GATA1s cells (42.6%) but were primarily MK-biased HPCs (**Figure 2H**), suggesting a maturation block. Thus, the effects of T21 and GATA1s compete in both erythropoiesis and megakaryopoiesis to give rise to cells with impaired lineage commitment, reminiscent of the aberrant MEP that underlies TAM.^15,17,18^

### GATA1s drives myelopoiesis in both euploid and T21 cells

The myeloid drive associated with GATA1s became apparent over time, particularly in euploid cells; by Day 11, 55.1% of euploid/GATA1s cells were myeloid compared to 9.3% of euploid/wtGATA1 (*p* <0.0001) (**Figure 2D**; **Table 1**). On Day 11, T21 was associated with increased myelopoiesis compared to euploid/wtGATA1 (T21/wtGATA1: 30.8%, T21/GATA1s: 35.8% vs. euploid/wtGATA1: 9.3%; *p* <0.0001 for both) (**Figure 2C-D**; **Table 1**). T21/wtGATA1 and T21/GATA1s cells had similar myeloid proportions (**Figure 2D**; **Table 1**), but interestingly, the GATA1s myeloid cells were predominantly in one cluster (**Figure 2C**).

### T21 and GATA1s independently and synergistically perturb gene regulation

GATA1s was associated with gene dysregulation in both euploid and T21 cells, although the number of DEGs varied with lineage and timepoint. On Day 7, GATA1s resulted in more DEGs in T21 cells compared to euploid in almost all populations (**Figure 3A**). Notably, there tended to be more DEGs identified between euploid and T21 cells with wtGATA1 than between euploid cells with wtGATA1 and GATA1s, suggesting that T21 and GATA1s each perturb gene regulation independently. On Day 11, GATA1s was associated with more DEGs in HPCs, MK-biased HPCs, and MKs in the euploid context, but more DEGs erythroid-biased HPCs, erythroid cells, and myeloid cells in the T21 context (**Figure 3B**). Comparison of Day 11 erythroid populations was limited since there were almost no euploid/GATA1s erythroid cells (**Table 1**).

**Figure 3.**
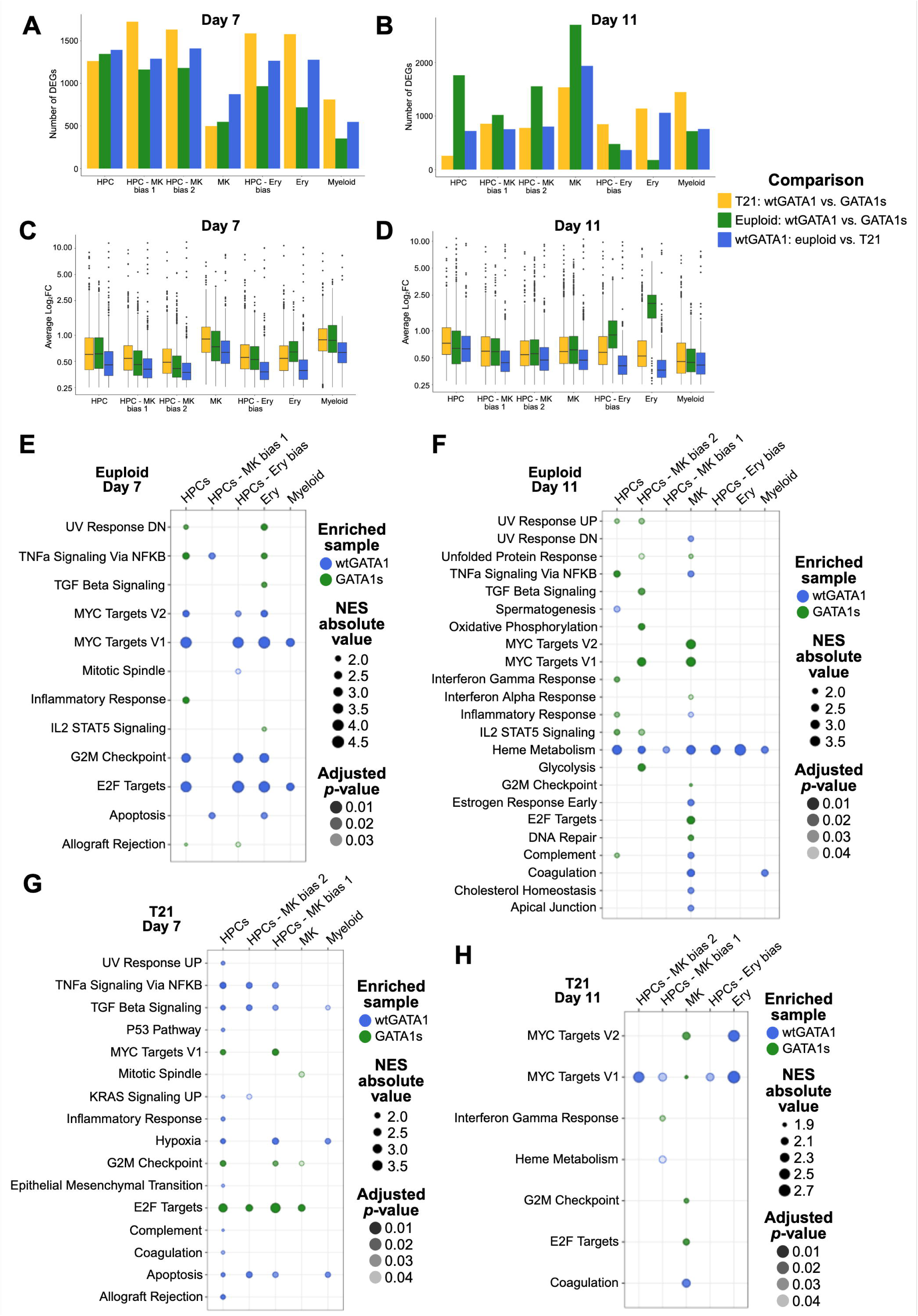
Differentially expressed genes and regulated pathways in HPCs generated from isogenic iPSCs differing by HSA21 and/or GATA1 status. A) Number of DEGs between wtGATA1 vs. GATA1s D7 and (B) D11 cells, organized by HSA21 status and lineage. C) Magnitude of log_2_FC of DEGs between wtGATA1 vs. GATA1s D7 and (D) D11 cells, organized by HSA21 status and lineage. E) GSEA of D7 euploid cells, (F) D11 euploid cells, (G) D7 T21 cells, and (H) D11 T21 cells, organized by lineage. Only gene sets represented in >1 lineage with NES ≥2.5 and adjusted *p*-value >0.02 are shown. NES = normalized enrichment score.

The mean log_2_FC of the DEGs tended to be highest between wtGATA1 and GATA1s cells with T21 and lowest between euploid and T21 cells with wtGATA1 (**Figure 3C-D**). Genes associated with the cell cycle, p53 network, histones, NF-κB network, globins, and platelet development and function were the most altered (**Figure 3E-H; Supplemental File 1**), consistent with known changes in hematopoietic and proliferative capacity with GATA1s.^38,62–65^ *MEG3*, a non-coding RNA that increases *p53* expression and activates p53 target genes,^66^ was strongly downregulated in T21, but not euploid, cells with GATA1s. Overall, many DEGs were recurrently identified (**Supplemental File 1**), and the timepoint– and lineage-specific differences further suggest isolated and synergistic effects of T21 and GATA1s on hematopoiesis.

### GATA1s exerts distinct effects on the cell cycle depending on lineage and timepoint

In the euploid context, GSEA on Day 7 cells revealed enriched pathways in all 7 clusters. MYC targets, G2M checkpoint genes, and E2F targets were enriched in wtGATA1 cells (**Figure 3E**), but by Day 11 these pathways were most enriched in MK-lineage cells with GATA1s and erythroid-lineage cells with wtGATA1 (**Figure 3F**). This may reflect a predominant effect of GATA1s in promoting proliferation during late-stage megakaryopoiesis at the expense of terminal maturation^67,63,64,68–74^ and the severe erythropoietic block in euploid/GATA1s cells, respectively. On Day 11, heme metabolism was enriched in wtGATA1 lineages, consistent with the severe erythropoietic block (**Figure 2C**).

In the T21 context, GSEA on Day 7 cells also revealed enrichment of MYC targets, G2M checkpoint genes, and E2F targets in HPCs, HPCs – MK bias 1 and 2, MKs, and myeloid cells, but in contrast to the euploid cells, the enrichment was associated with GATA1s (**Figure 3G**). Apoptosis was enriched in wtGATA1 cells. On Day 11, MYC targets were primarily enriched in wtGATA1 cells, but MYC targets, G2M checkpoint genes, and E2F targets were again enriched in GATA1s MKs (**Figure 3H**), suggesting that GATA1s enhances aberrant, proliferative megakaryopoiesis regardless of HSA21 background. The antiproliferative interferon-α response was uniquely enriched in T21/GATA1s, supporting prior findings that TAM cells overexpress interferon-α-responsive genes as a potential mechanism for spontaneous disease resolution.^75^ Taken together, the effect of GATA1s on cell cycle control appears to vary by HSA21 context and stage of cell differentiation.

### T21 and GATA1s alter the kinetics of lineage maturation

Since we observed altered MK, erythroid, and myeloid commitment associated with T21 and GATA1s, we performed trajectory inference on the integrated Day 7, 9, and 11 cells. This revealed the most mature cells in the UMAP periphery (**Figure 4A; Supplemental Figure 4A**). In the presence of wtGATA1, T21 was associated with impaired maturation of MKs and myeloid cells but enhanced maturation of erythroid cells when compared to euploid (**Figure 4B**). GATA1s was associated with delayed maturation of all lineages in both chromosomal contexts and tended to have a greater impact with T21. This was particularly notable in erythroid cells, in which GATA1s skewed T21 cells from the most to least mature of all genotypes.

Four terminal states—MK, erythroid, and myeloid 1 and 2—were inferred (**Figure 4C; Supplemental Figure 7B-C**) and validated using known marker genes (**Figure 4D**). The biological significance of the 2 myeloid states could not be distinguished with the available data. Latent time analysis of lineage-specific driver genes identified in this validation revealed the dynamic transcriptional programs associated with each terminal state (**Figure 4E**).

**Figure 4.**
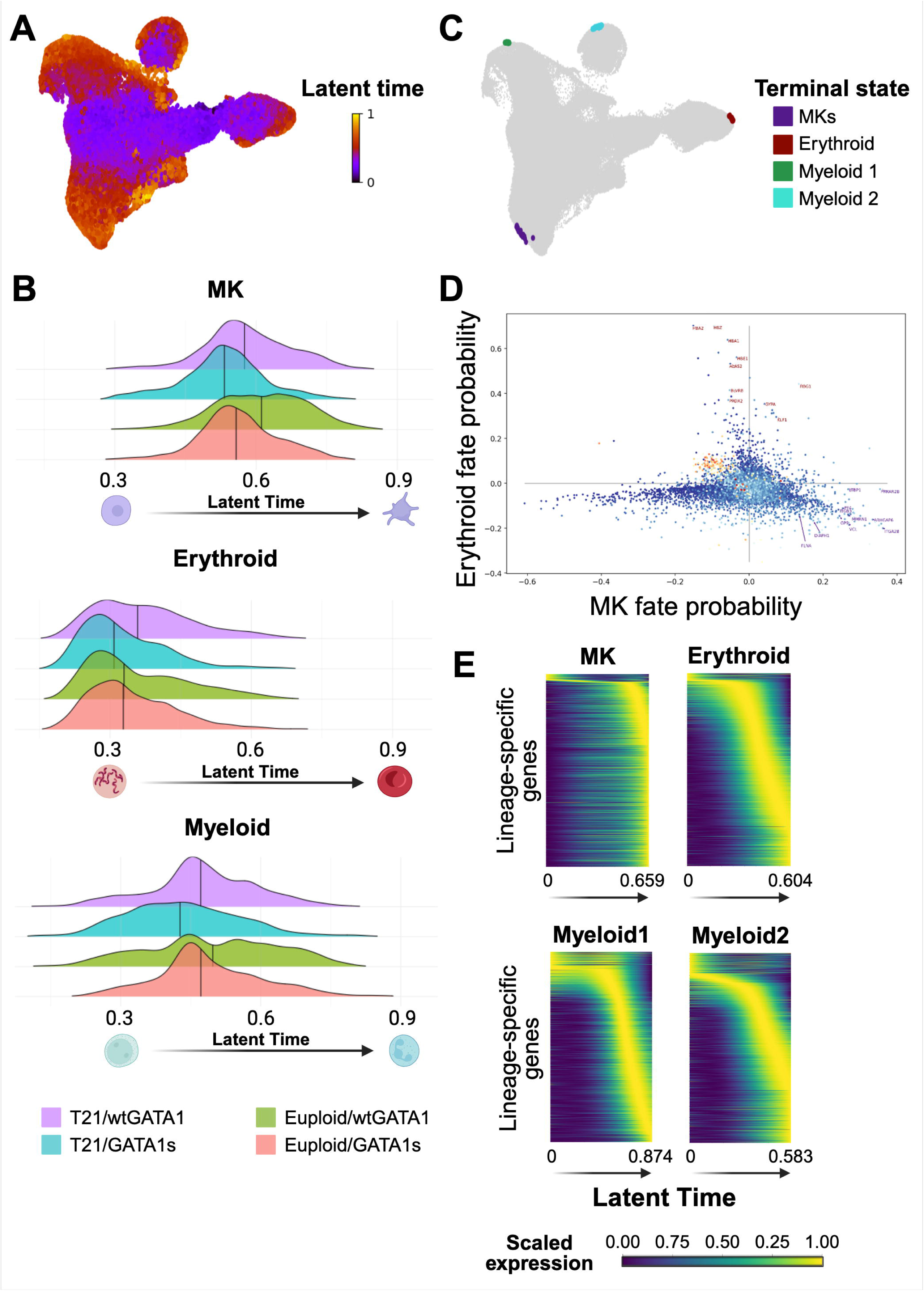
Trajectory analysis of HPCs generated from isogenic iPSCs differing by HSA21 and/or GATA1 status. A) Projection of latent time, as predicted by scVelo, indicating degree of differentiation and lineage maturation. 0 indicates least mature and 1 indicates most mature. B) Cell densities for MK, erythroid, and myeloid cells across all timepoints, separated by genotype. 0 indicates least mature and 1 indicates most mature. C) Terminal states inferred by latent time analysis. D) Sample plot of gene expression, correlated with fate probabilities, to identify driver genes for all trajectories. Upper left quadrant identifies driver genes for erythroid development and lower right quadrant identifies driver genes for MK development. E) Heatmaps of pseudotemporal gene expression across all samples for the 4 lineage-specific terminal states identified in (C). Genes ordered by peak expression time as cells progress towards terminal states. Latent time 0 indicates least mature and 1 indicates most mature.

### GATA1s perturbs lineage maturation through dysregulation of transcriptional programs

Genes that may drive the stalled maturation of T21/GATA1s cells were identified from lineage drivers that were also DEGs between wtGATA1 vs. GATA1s. The MK driver genes contained 4 subsets of expression patterns (**Figure 5A**). In euploid/wtGATA1 cells, the expression level of the 2 largest subsets increased with pseudotime, suggesting that these relate to maturation status (**Figure 5A purple/green**). Both T21/wtGATA1 and T21/GATA1s cells failed to upregulate expression of these gene subsets, consistent with impaired maturation. A small subset of genes was highly expressed in euploid/wtGATA1 cells throughout maturation, but inappropriately downregulated in T21/wtGATA1 cells and never expressed in T21/GATA1s cells (**Figure 5A yellow**). Finally, the expression levels of a fourth subset varied by genotype with no apparent relationship to pseudotime (**Figure 5A blue**). Overall, although euploid/GATA1s cells failed to upregulate MK driver genes to the extent of euploid/wtGATA1, T21/GATA1s strikingly had the lowest expression intensities. Taken together, this is consistent with our findings that GATA1s MKs exhibit a maturation block compared to wtGATA1, which is further exacerbated in the presence of T21 (**Figure 4C**). The MK driver genes were enriched for cell cycle pathways (MYC targets, G2M checkpoint, E2F targets, mitotic spindle); heme metabolism, which may reflect aberrant expression of an erythroid program; and mTORC1, which is required for normal *GATA1* expression during megakaryopoiesis and specifically maturation of neonatal MKs^76,77^ (**Figure 5B**).

**Figure 5.**
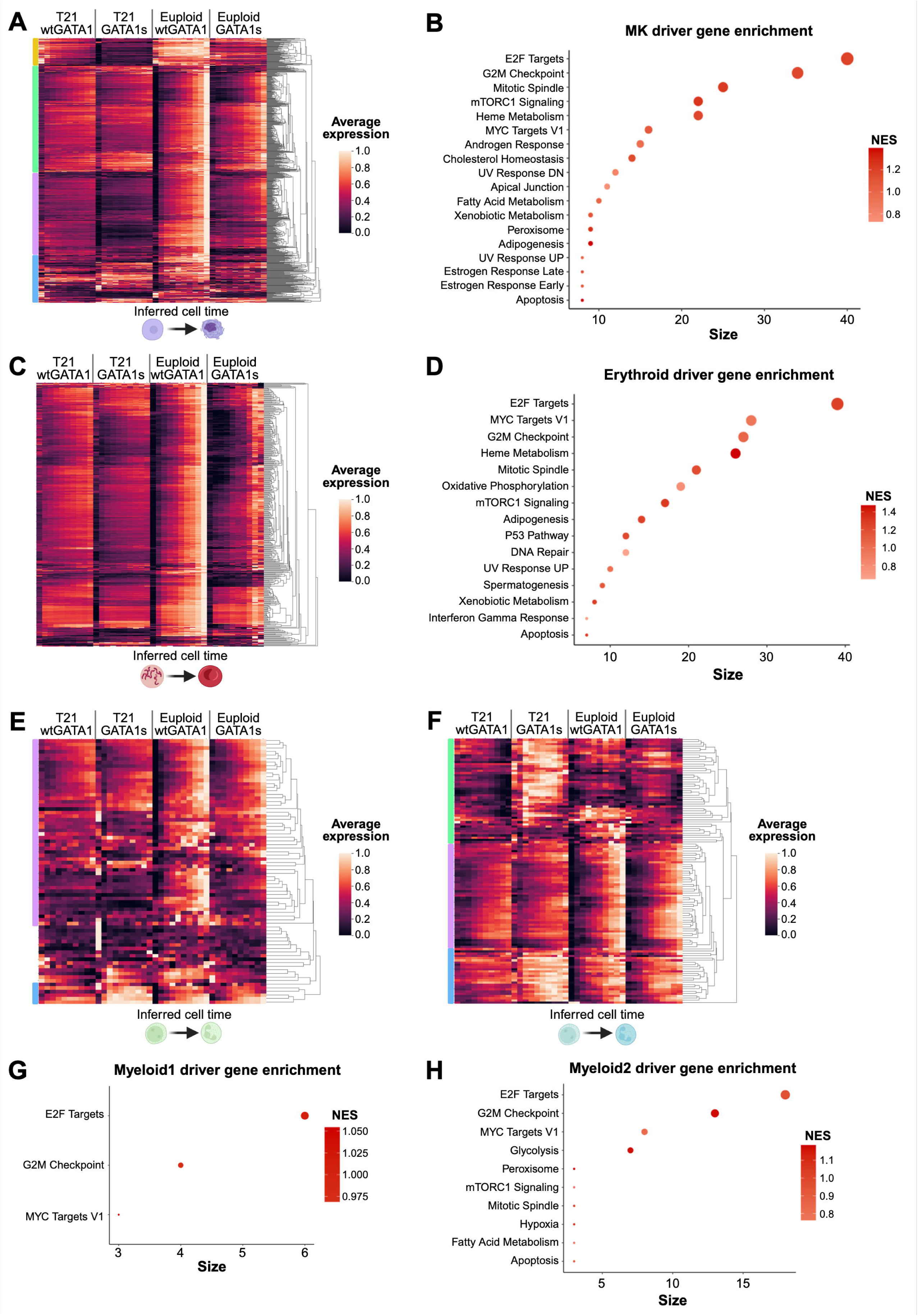
Analysis of driver genes associated with stalled lineage maturation of cells with T21 and/or GATA1s. Heatmaps showing developmental expression along pseudotime for driver genes identified between euploid vs. T21 and wtGATA1 vs. GATA1s for A) MKs C) erythroid cells E) myeloid 1 cells and F) myeloid 2 cells. Enrichment analysis of these driver genes for B) MKs D) erythroid cells G) myeloid 1 cells and H) myeloid 2 cells.

Erythroid driver genes were all progressively upregulated in euploid/wtGATA1 cells (**Figure 5C**). These genes were less uniformly regulated in euploid/GATA1s, with failure to upregulate some genes and premature upregulation of others. In contrast, in T21/wtGATA1 and T21/GATA1s cells, gene expression levels did not significantly change over pseudotime maturation, and T21/GATA1s cells again had the lowest expression intensity overall. These driver genes were enriched for cell cycle pathways and processes essential to erythroid differentiation including mTORC1 signaling,^78–80^ oxidative phosphorylation,^79^ and heme metabolism (**Figure 5D**).

Finally, the 2 myeloid terminal states (**Figure 4D**) displayed distinct patterns of gene regulation. Although the number of genes driving maturation towards myeloid 1 was limited, they tended to be progressively upregulated in euploid/wtGATA1 cells but only moderately upregulated in euploid/GATA1s cells and weakly with T21 (**Figure 5E purple**). A subset of genes was inappropriately upregulated throughout in T21/GATA1s (**Figure 5E blue**). The myeloid 2 driver genes segregated into three broad subsets. In euploid/wtGATA1 cells, 1 subset had expression apparently unrelated to pseudotime (**Figure 5F green**), and 2 subsets increased in expression intensity with pseudotime (**Figure 5F purple/blue**). Globally, the genes were insufficiently upregulated in euploid/GATA1s and T21/wtGATA1 cells. In contrast, 2 subsets of genes had inappropriately high expression levels with T21/GATA1s (**Figure 5F green/blue**). The driver genes for both myeloid states were enriched for E2F targets, G2M checkpoint genes and MYC targets (**Figure 5G-H**).

In all lineages, both T21 and GATA1s were associated with disruption of coordinated transcriptional programs, and GATA1s had distinct effects depending on euploid vs. T21 context. The recurrent enrichment of cell cycle genes suggests that the impaired maturation of T21/GATA1s cells is driven in part by an uncoupling between maturation and proliferation, consistent with a pro-leukemic phenotype.

## Discussion

Using a novel but well-characterized isogenic human iPSC system, we dissect clinically relevant effects of GATA1s on hematopoiesis in euploid and T21 HPCs as the basis of non-malignant cytopenias vs. TAM. Our isogenic euploid lines and single-cell approach build upon prior findings in iPSC-derived hematopoietic cells^37,38,40–42,70,81^ on how HSA21 copy number mediates the consequences of GATA1s on lineage commitment and maturation. In euploid cells, GATA1s nearly eliminates erythropoiesis and enhances MK and myeloid lineage commitment while slowing their maturation. In T21 cells, GATA1s competes with the erythroid drive of T21 and delays maturation of all 3 lineages. These findings are consistent with germline euploid *GATA1s* mutations typically causing anemia, MK dysplasia, and/or neutropenia,^21,82–84^ while T21 and GATA1s perturb the MEP and underlie both TAM and ML-DS. We demonstrate that the heterogeneity of HPCs is dynamically driven by genotype and that the effects of T21 and GATA1s are cell stage-specific.

While DS murine models have not fully recapitulated human TAM and ML-DS, some features overlap with our human cellular system. The bone marrow and spleen of adult DS mice contained increased erythroid cells^31^ and small, immature MKs,^31,33^ similar to the erythroid drive and delayed MK maturation in our T21/wtGATA1 cells. The proliferative, immature MKs generated from murine cells with *Gata1s*^13,69^ or deficient *Gata1*^64,67^ echo our findings of increased proliferation^38,70^ and impaired maturation of T21/GATA1s MKs. The early erythroid bias of our T21/GATA1s HPCs was surprising since they subsequently fail to differentiate into erythroid cells.^38,39^ However, other model systems provide evidence to support our findings. Arkoun et al found that T21/GATA1s iPSC-derived HPCs exhibited a similar incomplete suppression of erythropoiesis and a block in terminal MK maturation.^41^ Murine DS fetal livers with *Gata1s* contained increased erythroid progenitors although the mice were anemic,^31^ and GATA1s in CRISPR/Cas9-modified primary human euploid hematopoietic stem/progenitor cells caused a temporary accumulation of immature erythroblast-like cells.^85^

Our analyses suggest that temporal dysregulation of transcriptional programs underlies the hematopoietic perturbations, consistent with GATA1’s role as a transcription factor. The recurrent enrichment of MYC targets, E2F targets, and G2M checkpoint genes suggests a specific function for GATA1 in the cell cycle, particularly in MKs for which all 3 pathways were uniquely enriched on Day 11. Impaired *Myc* downregulation was found in murine *Gata1s* MKs,^13^ and conversely, sustained *c-MYC* overexpression in euploid iPSC-derived platelets was associated with decreased GATA1 levels and impaired platelet function.^86^ The N-terminus of GATA1 is required to block the interaction between E2F and its targets, which controls proliferation and terminal differentiation of HPCs into erythrocytes, MKs, and eosinophils.^70,87–89^ The enrichment in G2M checkpoint genes may reflect this dysregulation of E2F activity by GATA1s. Driver mutations in *ZBTB7A* and *IRX1* are found almost exclusively in ML-DS and upregulate the MYC/E2F pathway,^90^ suggesting that the dysregulation initiated by T21 and GATA1s may be requisite for rendering these “third-hit” mutations leukemogenic. Although we focused on how GATA1s perturbs euploid and T21 hematopoiesis, we also found enrichment of these cell-cycle pathways in T21/wtGATA1 MKs compared to euploid/wtGATA1 (data not shown), consistent with prior findings^29^ and suggesting that HSA21 copy number modulates GATA1’s interactions with MYC and E2F.

Our single-cell approach provides additional perspective on existing population-level studies. Comparisons of isogenic^91^ and non-isogenic^39^ T21/wtGATA1 and euploid/wtGATA1 iPSCs found that T21 enhances definitive HPC production and trilineage colony formation, but did not examine whether these HPCs would generate mature, lineage-committed cells. Our work suggests those HPCs may also be developmentally blocked, thus priming a population that gives rise to TAM. Prior studies with isogenic iPSCs found only minimal differences in the megakaryocytic skew of euploid and T21 cells with GATA1s,^39,40^ while our single-cell transcriptomes revealed significant differences depending on HSA21 copy number. Interrogation of genes such as *MEG3*, which had not been recognized by bulk RNA-seq of iPSC-derived T21 hematopoietic cells^42,70^ but was recently found to be downregulated in primary acute myeloid leukemia samples,^92^ may facilitate future studies for TAM and ML-DS therapies.

While the hematopoietic phenotypes associated with GATA1s and T21 have been studied extensively, isolating the underlying mechanisms has been challenging in part due to stage– and lineage-specific effects that may not be captured in prior human models. iPSCs provide a source of human multipotent HPCs that are otherwise difficult to obtain for study. Our single-cell approach reveals significant heterogeneity among stage-matched isogenic HPCs and that T21 and GATA1s synergistically alter lineage fate and enhance proliferation at the expense of maturation. Given the extensive number of known GATA1 targets and genome-wide driver genes identified here, our current findings are likely just one, albeit key, component of the hematopoietic perturbations that lead to TAM.

## Acknowledgements

The authors thank members of the CHOP Human Pluripotent Stem Cell Core Laboratory, Flow Cytometry Core, Center for Applied Genomics, Single Cell Technology Core, and High Throughput Sequencing Core for their expertise. This work was supported by the National Institutes of Health, National Heart, Lung, and Blood Institute T32 HL0007150 (KT), K99 HL156052 (CST), and R01 HL151260 (STC); American Society of Hematology Research Training Award for Fellows (KT); Doris Duke Charitable Foundation Physician Scientist Fellowship (KT); Mizuno Fund in Hematology (KT); and CHOP Distinguished Chair in Pediatrics (STC). Figures were compiled and edited in BioRender.

## Author Contributions

KT designed research studies, performed experiments, acquired data, interpreted the data, and wrote the manuscript. EW and DS analyzed data and edited the manuscript. SSK, YTS, and AG performed experiments. DLF and CST interpreted the data and edited the manuscript. STC designed research studies, interpreted the data, and wrote the manuscript.

## Disclosure of Conflicts of Interest

The authors have no conflicts of interest to declare.

## Supplemental Methods

### Western blot

Cells were lysed in Pierce RIPA Buffer (ThermoFisher) with 1% Halt Protease Inhibitor and Phosphatase Inhibitor Cocktails (ThermoScientific). Protein contents of lysates were normalized using the Pierce BCA Protein Assay Kit (ThermoFisher) according to the manufacturer’s instructions, fractionated on Bolt 4-12% Bis-Tris gradient gels (Invitrogen), and transferred to 0.2 μm PVDF membranes (GE Healthcare). SuperSignal^TM^ West Femto Maximum Sensitivity Substrate was used for imaging on a KwikQuant Imager (Kindle Biosciences). Primary and secondary antibodies are shown in **Supplemental Table 2**.

### Hematopoietic differentiation of iPSCs

iPSCs were cultured and differentiated into hematopoietic cells by embryoid body (EB) formation using modifications on a protocol previously described by our laboratory.^37,38,47,48^ On the initial day of differentiation (Day-2), iPSCs were incubated with Accutase (ThermoFisher Scientific) at 37°C x 2 minutes, then washed with phosphate-buffered saline (PBS) x 2 to remove the mouse embryonic fibroblast feeder layer. The iPSCs were resuspended in human embryonic stem media with 10 ng/mL of bFGF (beta fibroblast growth factor, Invitrogen) (HES-10) and ROCK inhibitor (10 μM final concentration, Cayman Chemical) as aggregates of 5-10 cells each at a density of 1.5-2 x 10^6^ cells/2mL per well in 6-well ultra-low attachment plates (Corning).

Cells were maintained at 37 °C, 5% CO_2_, and 5% O_2_ on a rotary shaker at 100 rotations per minute. On Days –1 and 0, EBs were isolated by gravity x 20 minutes. EBs were resuspended in HES-10 on Day –1, and in StemPro-34 (Invitrogen) with glutamine (1%, Gibco), ascorbic acid (50 ng/mL, Sigma-Aldrich), monothioglycerol (3 μL/mL, Sigma-Aldrich), BMP-4 (bone morphogenic protein 4, 25 ng/mL, Invitrogen) and VEGF (vascular endothelial growth factor, 10 ng/mL, Invitrogen) on Day 0 prior to redistribution into the 6-well plates. EBs were subsequently cultured in multi-cytokine differentiation media to induce mesoderm and hematoendothelial specification as previously described (**Figure 1C**).^37,38^

### Isolation of hematopoietic progenitors

On Day 7, HPCs were collected from disaggregated EBs and the supernatant. EBs were isolated by centrifugation at 600 rpm x 1 minute, then chemically disaggregated in collagenase B (Sigma-Aldrich) at 37 °C x 60 minutes and then 0.25% trypsin-EDTA (Gibco) at room temperature x 2 minutes followed by trypsin inactivation with 50% fetal bovine serum (FBS, GeminiBio) in IMDM (Iscove’s Modified Dulbecco’s Medium, Corning). EBs were then mechanically disaggregated by passing through a 20G needle/5mL syringe and filtered through a 40 μm cell strainer to remove stromal debris to isolate the HPCs. HPCs were isolated from the supernatant by additional centrifugation at 1200 rpm x 5 minutes. HPCs from both sources were combined and incubated with fluorescently-conjugated antibodies against CD41a and CD235a (**Supplemental Table 3**) in FACS buffer (0.1% bovine serum albumin (BSA), 0.0002% NaN_3_, 0.005 M ethylenediaminetetraacetic acid (EDTA) in PBS) at 4 °C x 20 minutes.

**Supplemental Table 1.**
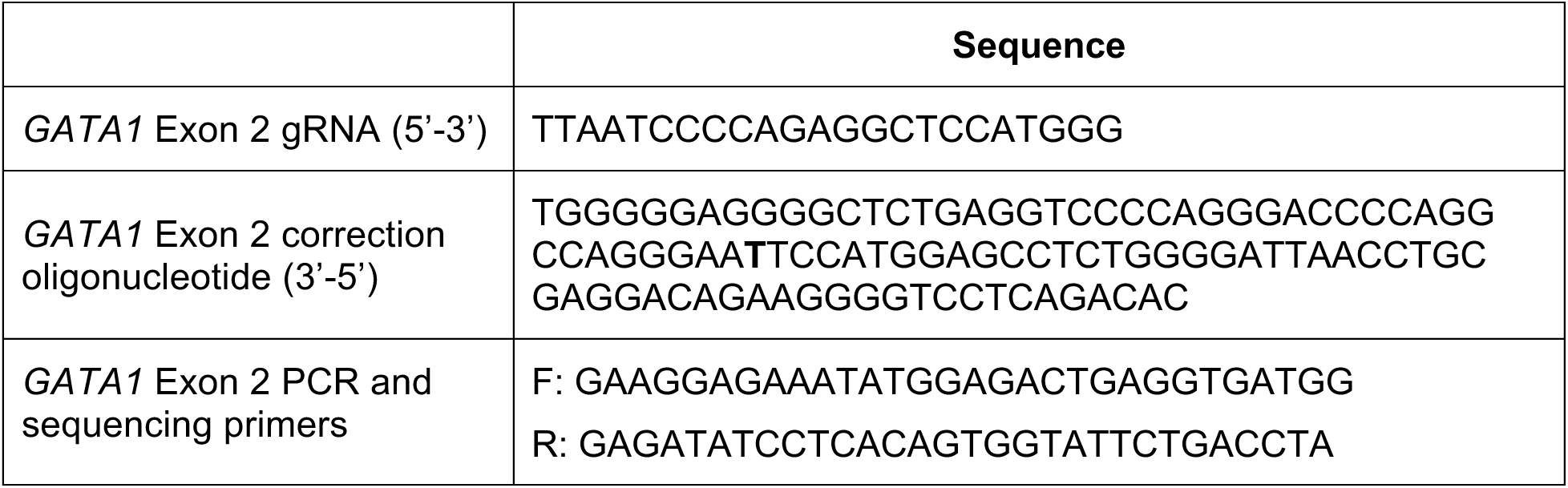
CRISPR/Cas9 targeting guide RNAs, oligonucleotides for homologous recombination, and PCR and sequencing primers.

**Supplemental Table 2.** Antibodies for Western blot.

| **Antibody** | **Primary/Secondary** | **Manufacturer** | **Catalog #** | **Dilution** |
| --- | --- | --- | --- | --- |
| GATA1 | Primary | Cell Signaling Technology | 4589 | 1:1000 |
| GAPDH | Primary | Abcam | ab181602 | 1:10000 |
| Rabbit IgG<br>(HRP-conjugated) | Secondary | Cell Signaling Technology | 7074 | 1:10000 |

**Supplemental Table 3.** Antibodies for flow cytometry.

| **Antibody** | **Fluorophore** | **Manufacturer** | **Catalog #** | **Dilution** |
| --- | --- | --- | --- | --- |
| CD18 | FITC | Fisher Scientific | 555923 | 1:20 |
| CD235a | APC | BD Biosciences | 551336 | 1:5000 |
| CD34 | APC | BioLegend | 561209 | 1:100 |
| CD41a | PE/Cy7 | BioLegend | 303718 | 1:800 |
| CD42a | FITC | BD Biosciences | 558818 | 1:20 |
| CD45 | APC | BD Biosciences | 55485 | 1:20 |

**Supplemental Table 4.** Mitochondrial RNA content of each sample.

| Sample |  | Mitochondrial RNA content |
| --- | --- | --- |
| Day 7 | Euploid/wtGATA1 | 25% |
|  | Euploid/GATA1s | 20% |
|  | T21/wtGATA1 | 15% |
|  | T21/GATA1s | 15% |
| Day 9 | Euploid/wtGATA1 | 15% |
|  | Euploid/GATA1s | 15% |
|  | T21/wtGATA1 | 15% |
|  | T21/GATA1s | 25% |
| Day 11 | Euploid/wtGATA1 | 15% |
|  | Euploid/GATA1s | 15% |
|  | T21/wtGATA1 | 15% |
|  | T21/GATA1s | 15% |

**Supplemental Table 5.** Manually selected hematopoietic genes for cluster annotation.

| Gene | Associated lineage | Gene | Associated lineage | Gene | Associated lineage |
| --- | --- | --- | --- | --- | --- |
| *ACHE* | Ery | *CNGB1* | MK | *GATA3* | HPC |
| *ACLY* | MK | *COPA* | MK | *GFI1* | MK |
| *ACSL1* | Myeloid | *CREB1* | MK | *GFI1B* | MK |
| *ACTB* | Housekeeping | *CSF1R* | Myeloid | *GLO1* | Ery |
| *ADD1* | Ery | *CSF2RA* | Myeloid | *GP1BA* | MK |
| *ADD2* | Ery | *CSF3R* | Myeloid | *GP9* | MK |
| *ALAS2* | Ery | *CSTA* | Myeloid | *GPD2* | MK |
| *ALB* | MK | *CST3* | Ery | *GPI* | MK |
| *ALDH1A1* | Ery | *CTSE* | Ery | *GSR* | Ery |
| *ANGPT1* | HPC | *CXCL8* | Myeloid | *GTF2H4* | MK |
| *ANK1* | MK | *CYB5A* | Ery | *GYPA* | Ery |
| *ARHGAP6* | MK | *CYB5R3* | Ery | *HBA1* | Ery |
| *ARHGAP15* | HPC | *CYBB* | Myeloid | *HBA2* | Ery |
| *ATP2B1* | Ery | *DAAM1* | MK | *HBB* | Ery |
| *ATP2B4* | Ery | *DAAM2* | MK | *HBE1* | Ery |
| *AZU1* | Myeloid | *DCTN1* | MK | *HBG1* | Ery |
| *BACH1* | Ery | *DDI1* | MK | *HBZ* | Ery |
| *BBS1* | Ery | *DIAPH1* | MK | *HDHD1* | Ery |
| *BCL11A* | Ery | *DMTN* | Ery | *HK1* | MK |
| *BIN2* | MK | *DYNC1H1* | MK | *HMBS* | Ery |
| *BLVRB* | Ery | *DYRK1A* | MK | *HMHA1* | MK |
| *BMI1* | Ery | *EGR2* | Myeloid | *HOXA10* | HPC |
| *CASS4* | MK | *EHD1* | MK | *HOXA9* | HPC |
| *CAT* | Ery | *ENPP1* | MK | *HSP90AA1* | MK |
| *CCDC181* | MK | *EPB41* | Ery | *HSP90B1* | MK |
| *CCL3* | Myeloid | *EPB42* | Ery | *IFI16* | Myeloid |
| *CCL4* | Myeloid | *EPOR* | Ery | *IKZF1* | Myeloid |
| *CCT3* | MK | *ERG* | MK | *IKZF2* | Myeloid |
| *CD34* | HPC | *ETS2* | MK | *IL8* | Myeloid |
| *CD44* | Ery | *EVI1* | HPC | *INF2* | Myeloid |
| *CD47* | Ery | *F10* | MK | *IQGAP2* | HPC |
| *CD58* | Ery | *F5* | MK | *IRAK3* | Myeloid |
| *CD74* | Myeloid | *FGA* | MK | *IRF1* | Myeloid |
| *CEBPA* | Myeloid | *FGB* | MK | *IRF7* | Myeloid |
| *CEBPB* | Myeloid | *FLI1* | HPC/MK | *ITGA1* | MK |
| *CEBPG* | Myeloid | *FLNA* | MK | *ITGA4* | HPC |
| *CFAP36* | MK | *FOXO3* | HPC | *ITGA2B* | MK |
| *CFH* | Myeloid | *GABPA* | HPC | *ITGB3* | MK |
| *CLC* | Myeloid | *GAPDH* | Housekeeping | *JAK3* | Myeloid |
| *CLTC* | MK | *GATA1* | Ery | *KIF16B* | MK |
| *CREM* | MK | *GATA2* | HPC | *KIT* | Ery |
| *KLF1* | Ery | *PRDX6* | Ery | *TRIM10* | Myeloid |
| *LANCL1* | Ery | *PRKAR2B* | MK | *TSTA3* | Ery |
| *LDB1* | Ery | *PRG2* | Myeloid | *UBE2O* | MK |
| *LGMN* | Myeloid | *PRG3* | Myeloid | *UBR4* | MK |
| *LTBP1* | MK | *PRTN3* | Myeloid | *UGGT1* | MK |
| *LTF* | MK | *PSPC1* | MK | *UGGT2* | MK |
| *LYL1* | Myeloid | *PYGB* | MK | *UNC13D* | MK |
| *MED12L* | MK | *PYGL* | MK | *UTRN* | MK |
| *MEIS1* | HPC | *RBM47* | Myeloid | *VCL* | MK |
| *MMRN1* | MK | *RNF220* | HPC | *VCP* | MK |
| *MPL* | Myeloid | *RCAN1* | Myeloid | *VDR* | Myeloid |
| *MPO* | Myeloid | *RUNX1* | HPC/MK | *VIM* | Myeloid |
| *MPP1* | Ery | *S100A8* | Myeloid | *VPS13A* | MK |
| *MRC1* | Myeloid | *S100A9* | Myeloid | *VWF* | MK |
| *MS4A3* | Myeloid | *SAMHD1* | Myeloid | *WDFY3* | MK |
| *MT1E* | Ery | *SDHA* | Housekeeping | *WDR1* | MK |
| *MYB* | HPC | *SELP* | MK | *WDR44* | MK |
| *MYC* | HPC | *SKP1* | Ery | *WNT10A* | HPC |
| *MYO18A* | MK | *SLC12A1* | MK | *WNT8B* | MK |
| *NAB2* | Myeloid | *SLC4A1* | Ery | *YARS* | MK |
| *NBEAL1* | MK | *SMAD1* | HPC | *ZC3H12C* | Myeloid |
| *NBEAL2* | MK | *SMC3* | MK | *ZFPM1* | MK |
| *NCAM1* | Myeloid | *SPINK2* | HPC | *ZNF559* | MK |
| *NCF1* | MK | *SON* | HPC | *ZSCAN12* | MK |
| *NEXN* | MK | *SOX17* | HPC |  |  |
| *NFE2* | HPC | *SOX4* | HPC |  |  |
| *NFX1* | HPC | *SOX6* | Ery |  |  |
| *NRP1* | Myeloid | *SP1* | Myeloid |  |  |
| *P4HB* | MK | *SPTA1* | MK |  |  |
| *PBX1* | HPC | *SPTB* | MK |  |  |
| *PCMT1* | Ery | *SPTBN2* | MK |  |  |
| *PDE4D* | Myeloid | *SPTBN4* | MK |  |  |
| *PDIA4* | MK | *STAT2* | Myeloid |  |  |
| *PDZD3* | MK | *STAT3* | Myeloid |  |  |
| *PF4* | MK | *STIP1* | MK |  |  |
| *PF4V1* | MK | *STOM* | Ery |  |  |
| *PGK1* | Ery | *SYTL4* | MK |  |  |
| *PIEZO2* | MK | *TAF1B* | MK |  |  |
| *PLSCR1* | Ery | *TAL1* | HPC |  |  |
| *PPBP* | MK | *TFEC* | HPC |  |  |
| *PPIA* | HPC | *TMC1* | MK |  |  |
| *PRDX2* | Ery | *TPP2* | MK |  |  |

## Supplemental Figure Legends

**Supplemental Figure 1.**
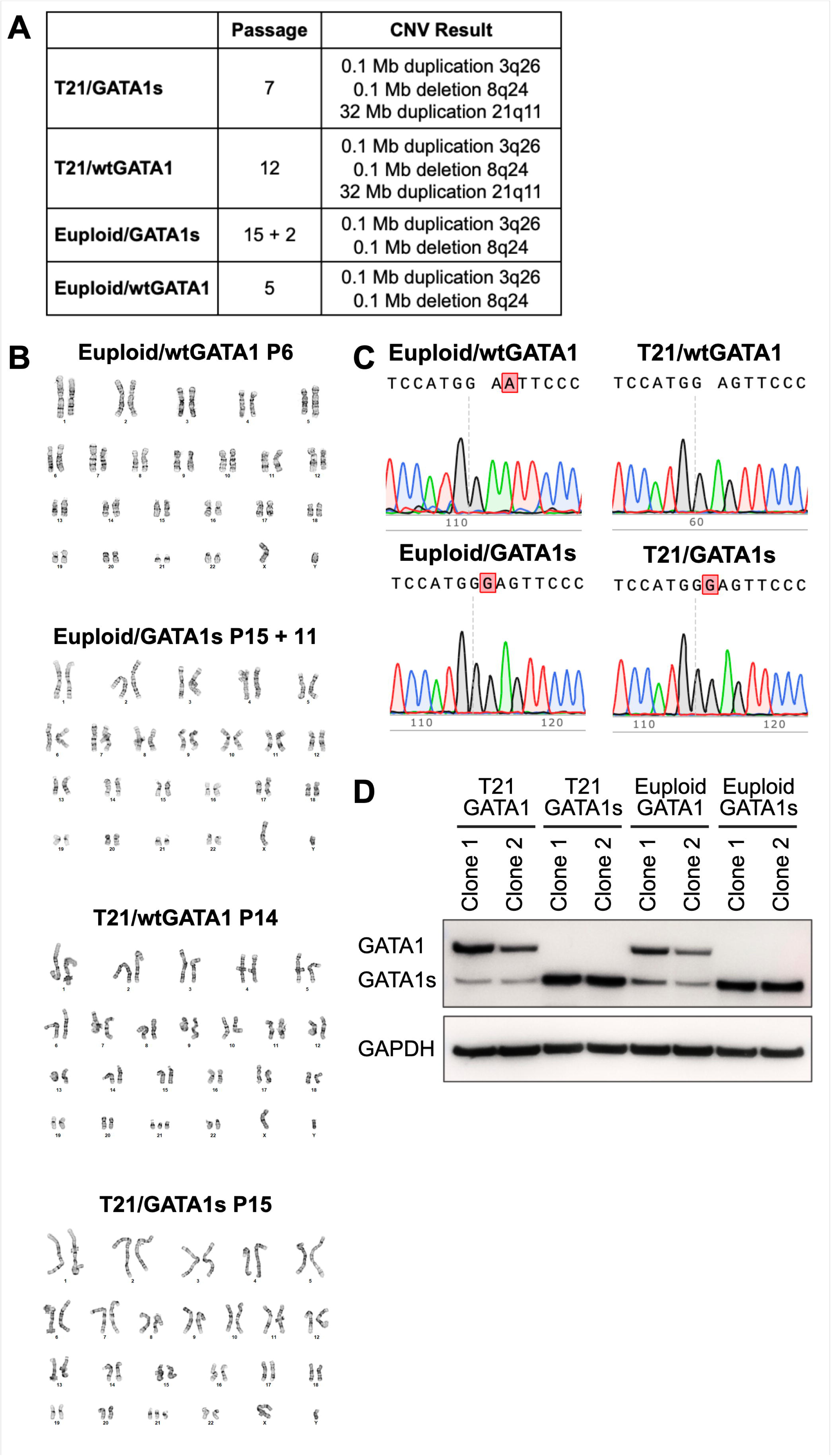
Characterization of isogenic iPSCs derived from single individual with TAM. A) Copy number variation analysis of iPSCs. B) Karyotypes of iPSCs indicating 46,XY or 47,XY+21. C) Sanger sequencing of iPSCs showing the patient-specific c.4_5insG mutation in iPSC lines with GATA1s. Additional c.6G>A mutation in euploid/wtGATA1 line is silent mutation introduced during CRISPR/Cas9 gene targeting. D) Western blot showing expression of GATA1 and GATA1s in iPSC-derived MKs. Images were acquired from one gel using KwikQuant Imager and processed in Adobe Photoshop.

**Supplemental Figure 2.**
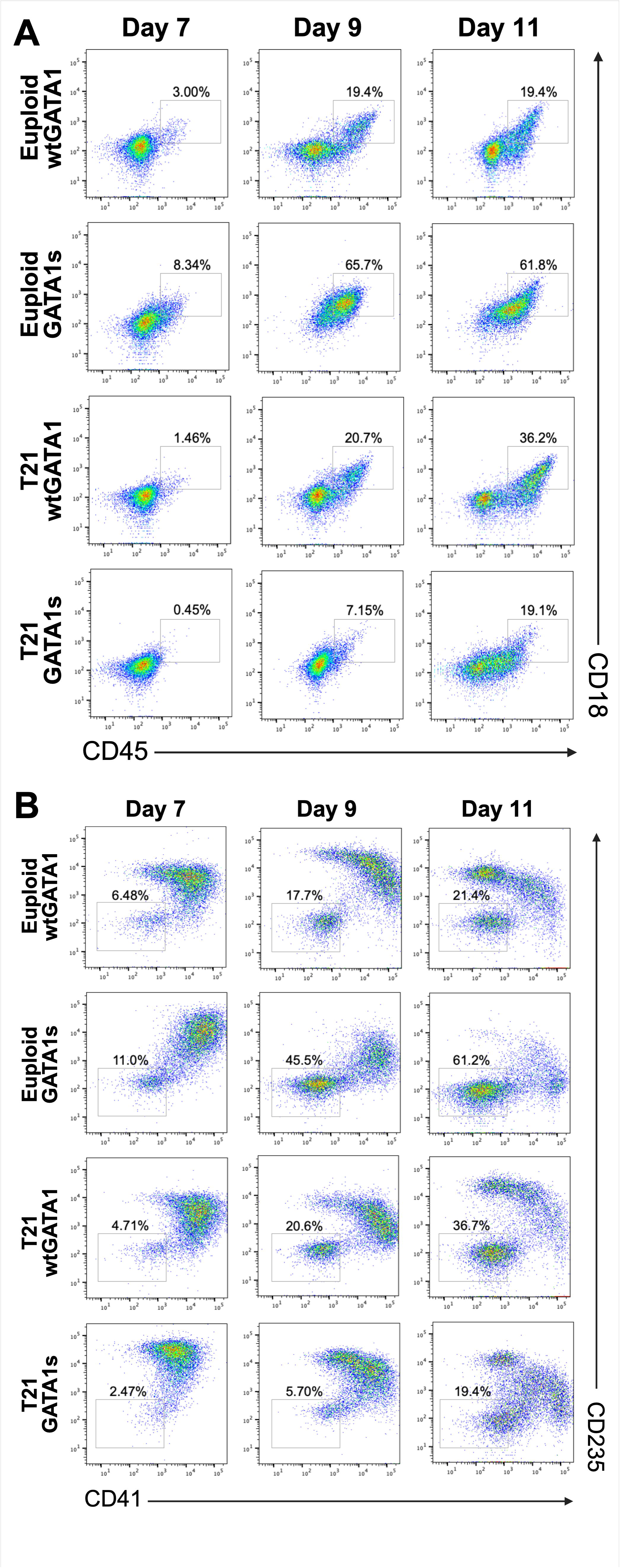
Flow cytometry plots of Day 7, Day 9, and Day 11 hematopoietic progenitors differing only by HSA21 and/or GATA1 status. A) CD45^+^/CD18^+^ indicates myeloid bias, with percentages approximating the CD41^-^/CD235^-^ population on corresponding flow cytometry plot in (B).

**Supplemental Figure 3.**
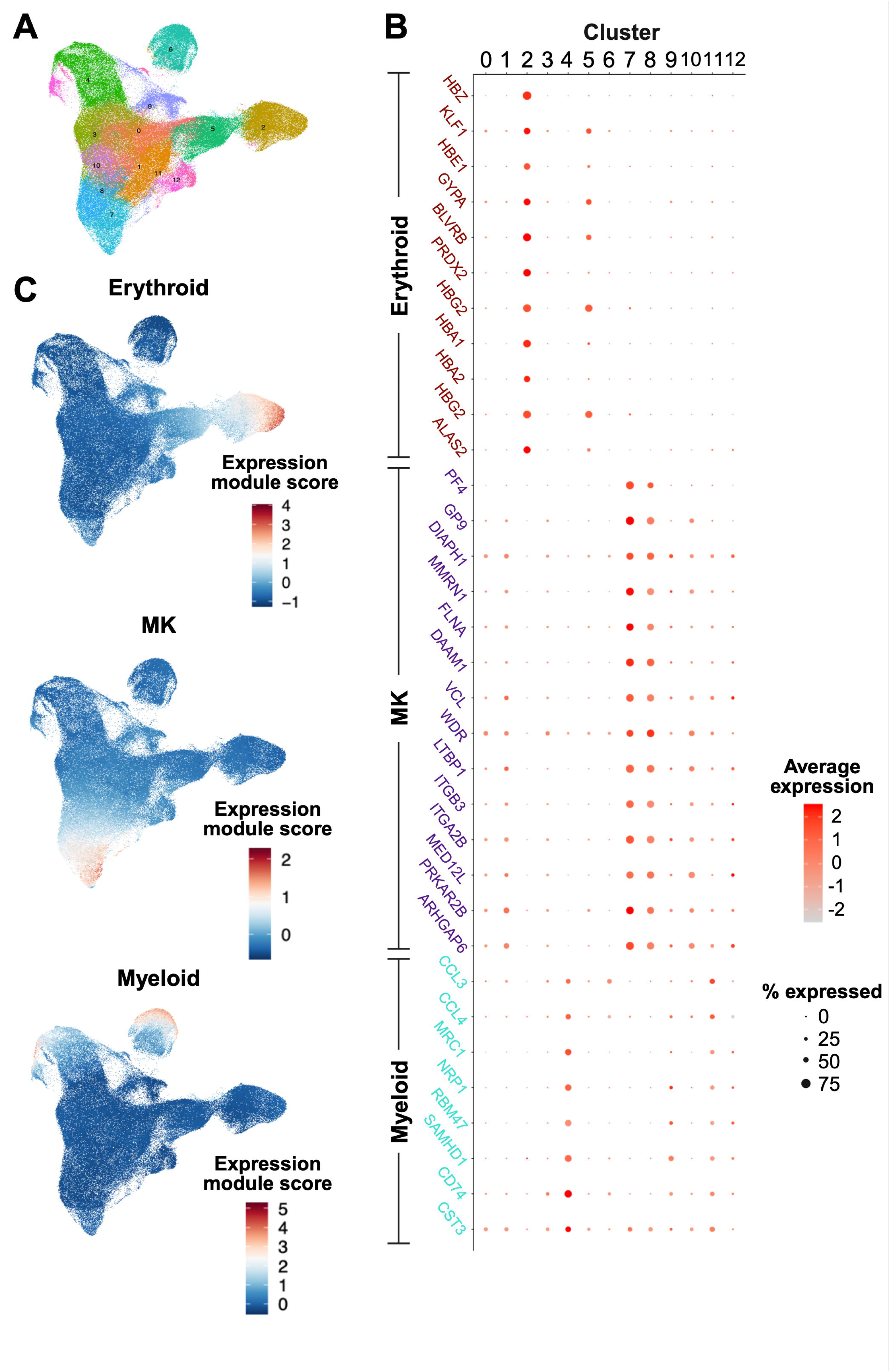
Annotation of integrated UMAP of euploid/wtGATA1, euploid/GATA1s, T21/wtGATA1, and T21/GATA1s hematopoietic progenitor cells (HPCs) collected on Day 7, Day 9, and Day 11. A) Unannotated integrated UMAP showing 13 distinct clusters. B) Expression of representative key hematopoietic genes in each cluster, distinguishing clusters 2 and 5 as likely erythroid lineage, clusters 0, 1, 3, 7, 8, and 10 as likely MK lineage, and 4 and 6 as likely myeloid lineage. C) Lineage-enriched (erythroid, MK, and myeloid) UMAP regions based on Seurat expression module scores for all lineage-specific genes. Higher values correspond to higher expression levels of lineage-specific genes.

**Supplemental Figure 4.**
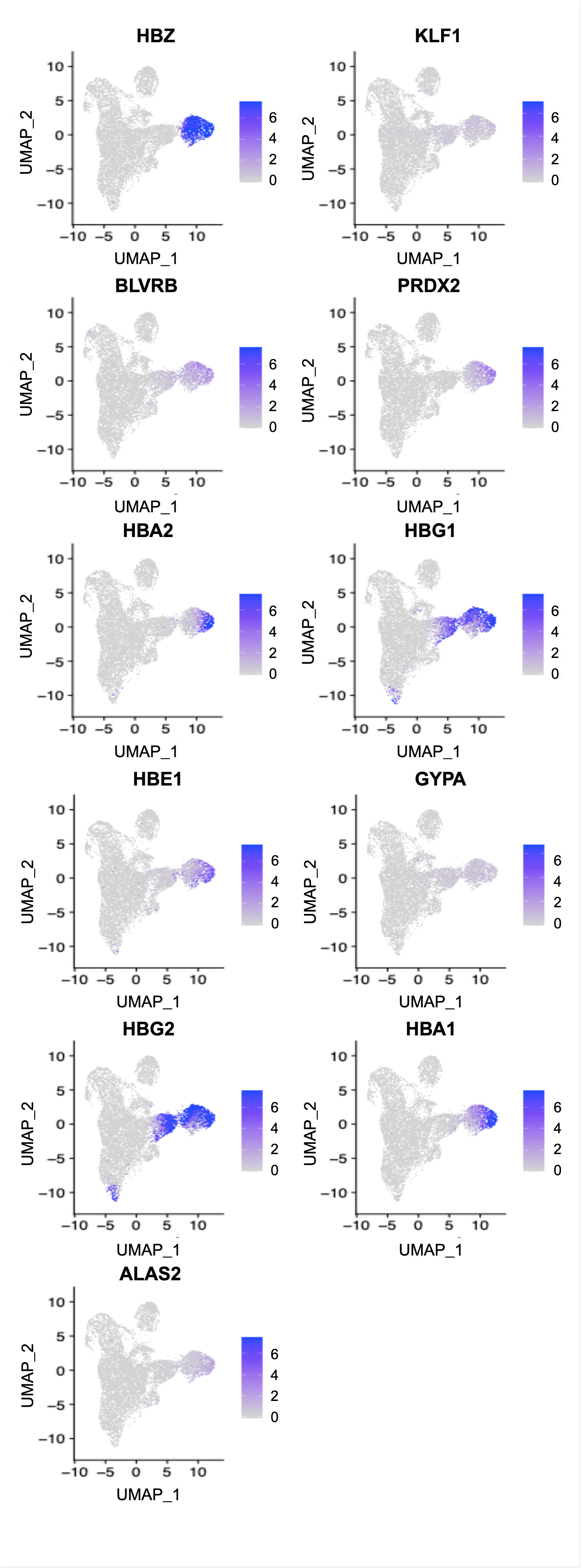
Key erythroid-specific genes with distinct expression between UMAP regions. Expression levels identify clusters 2 (erythroid) and 5 (HPC – erythroid bias as likely erythroid lineage. Higher values correspond to higher expression levels.

**Supplemental Figure 5.**
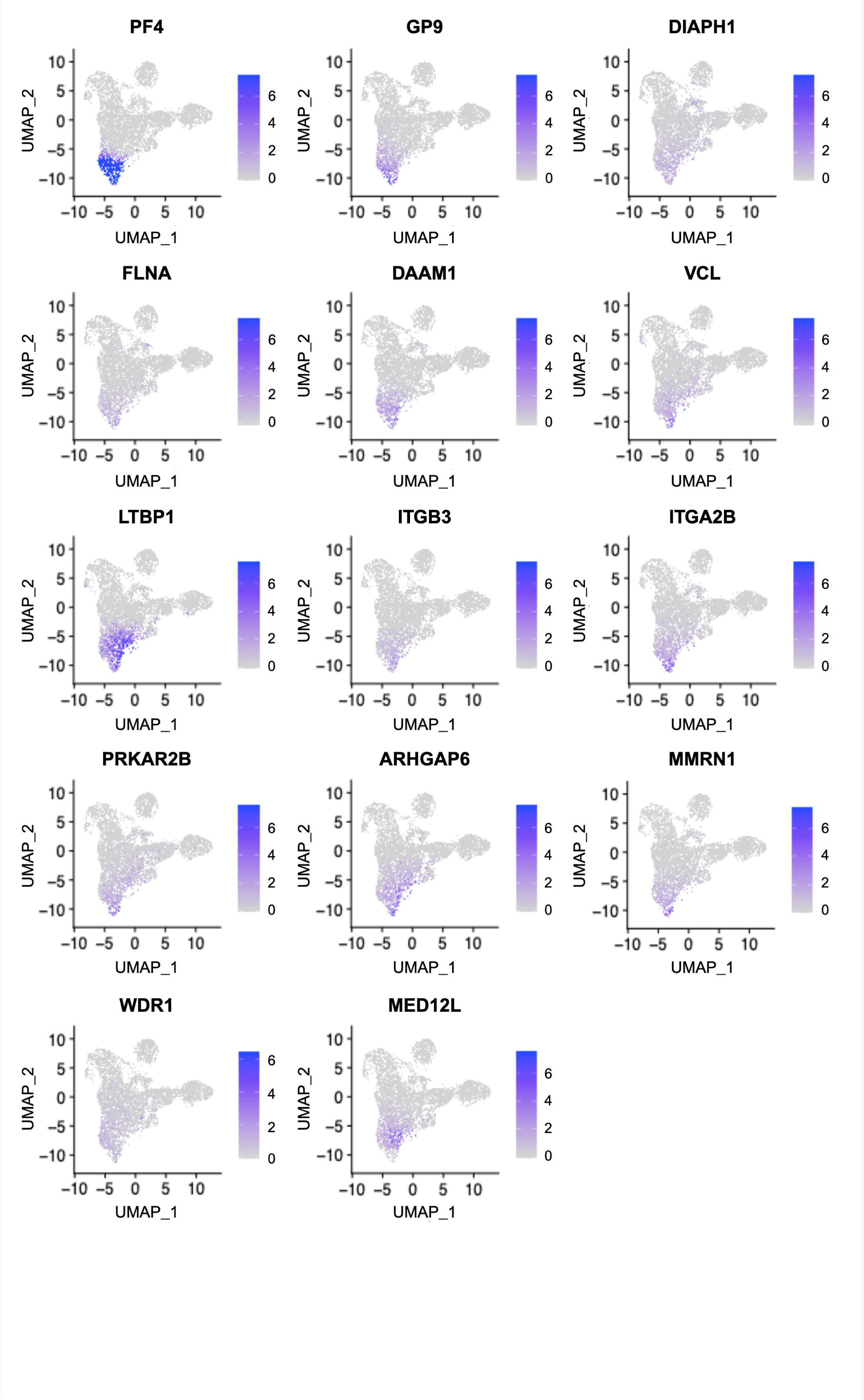
Key megakaryocyte (MK)-specific genes with distinct expression between UMAP regions. Expression levels identify clusters 7 and 8 (MK), 1 and 10 (HPC – MK bias 1), and 0 and 3 (HPC – MK bias 2) as likely MK lineage. Higher values correspond to higher expression levels.

**Supplemental Figure 6.**
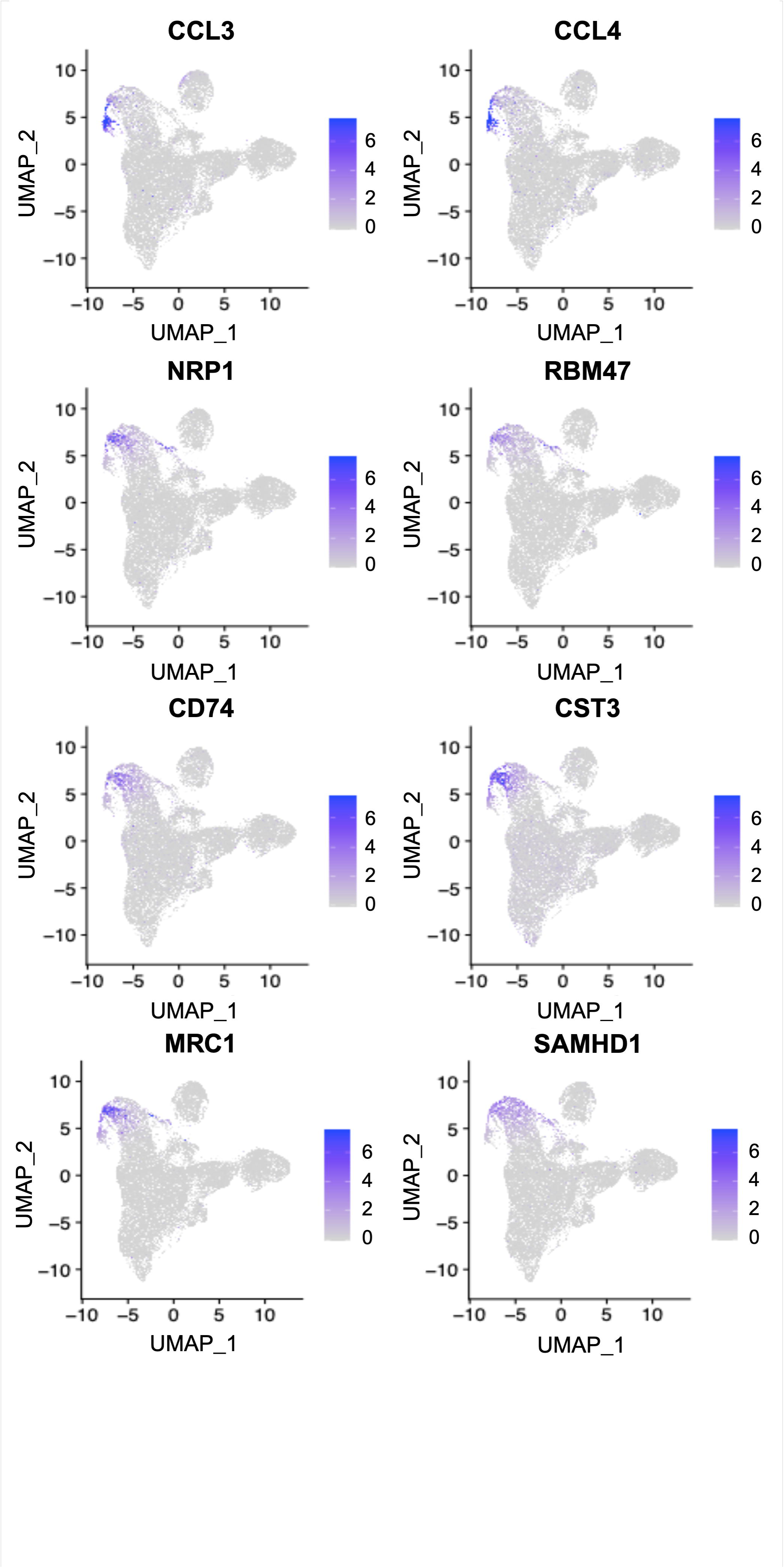
Key myeloid-specific genes with distinct expression between UMAP regions. Expression levels identify clusters 4 and 6 as likely myeloid lineage. Higher values correspond to higher expression levels.

**Supplemental Figure 7.**
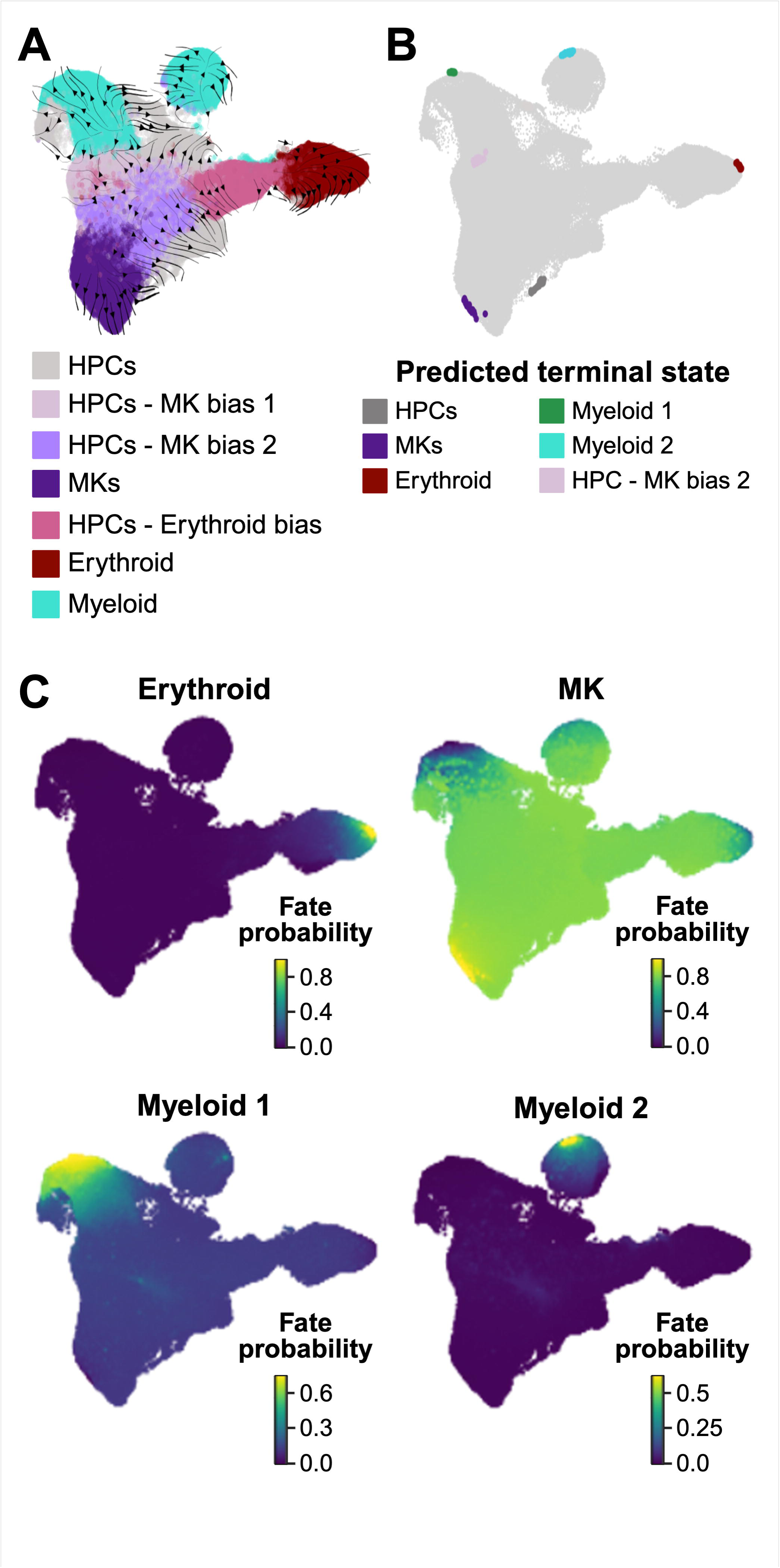
Pseudotime projections of differentiation into terminal lineages. A) UMAP of pseudotime velocity analysis, with arrows indicating direction of transition towards cell states. Arrows indicate direction of cell state transition. B) Predicted terminal states from which the four terminal states (erythroid, MKs, myeloid 1, and myeloid 2) were selected. C) Cell fate probabilities towards selected terminal states (erythroid, MKs, myeloid 1, and myeloid 2) as predicted by CellRank-CytoTRACE. 0 indicates least likely and 1 indicates most likely.

**Supplemental File 1.** Full list of DEGs on Days 7 and 11, organized by gene name. Analyses were performed separately for each lineage/cell type, and DEGs were identified between wtGATA1 vs. GATA1s cells within each condition (euploid or T21).

## Notes

### Competing Interest Statement

The authors have declared no competing interest.

### Summary of Updates

This revised version of the manuscript includes additional data on cluster annotation, more robust interpretations of trajectory inference, and a more extensive discussion including references to works published since the initial preprint.

